# Neural correlates of image memorability: Combining large-scale 7T fMRI with machine learning-based predictions

**DOI:** 10.1101/2025.04.06.647493

**Authors:** V. Mäki-Marttunen, T. Hagen, T. Espeseth

## Abstract

We combine ultra-high-resolution fMRI with machine learning predictions to investigate the neural correlates of image memorability. Using ViTMem, a visual transformer-based tool, we estimated memorability scores for over 70,000 natural scenes from the NSD dataset. Our analyses reveal that higher memorability scores correlate with increased activation in ventral visual regions (fusiform gyrus, middle occipital cortex) and medial temporal areas (hippocampus, amygdala). Notably, we also observed significant engagement in fronto-parietal regions, indicating a broader network involved in modulating memory encoding. These novel findings demonstrate that ViTMem not only reliably predicts stimulus memorability but also provides a powerful framework for scalable investigations into the neural mechanisms underlying perceptual and mnemonic processes.

## Introduction

Image memorability, the extent to which certain images are consistently remembered or forgotten by individuals (Kramer et al. 2023), concerns how humans focus on and process new visual information. An important property of visual stimuli is that their memorability, rather than being a binary distinction between remembered and forgotten images, can be quantified on a scale, referred to as a memorability score. Prior research using functional magnetic resonance imaging (fMRI) has provided insights into the neural correlates of image memorability, revealing that specific brain regions, particularly higher visual circuits and medial temporal lobe regions, play vital roles in encoding memorable visual stimuli (Bainbridge et al., 2017; Bainbridge & Rissman 2018; Lahner et al. 2024). In contrast, fronto-parietal brain regions have been found to be associated with successful memory retrieval, but not with stimulus memorability (Bainbridge et al. 2017; Bainbridge & Rissman, 2018). This suggests that the neural substrates of image memorability may be restricted to perceptual and mnemonic processes in the ventral visual pathway and medial temporal lobe. However, these findings are based on relatively few studies with limited trial numbers and stimulus sets, leaving open questions about the extent and stability of the identified correlates of image memorability. To advance our understanding of the neural basis of image memorability beyond these limitations, we developed a novel approach that combines artificial neural networks (ANNs) trained to predict image memorability with large-scale fMRI datasets. In this study, we utilized a publicly available ultra-high-resolution fMRI dataset with a large number of trials and unique stimuli. We estimated the memorability of each stimulus using ViTMem, our visual transformer-based image memorability prediction tool (Hagen & Espeseth, 2023). We hypothesized that the analysis would reveal clear evidence of sensitivity to image memorability in the ventral visual cortex and medial temporal lobe regions, but not in fronto-parietal cortical areas.

## Methods

### Sample

We used the NSD dataset available at https://naturalscenesdataset.org (Allen et al. 2022). The dataset consisted of 7T fMRI acquisitions performed during the presentation of ~70,500 unique, coloured natural scenes. Images were selected from the Microsoft Common Objects in Context (COCO) dataset (Lin et al. 2015), which contains a broad selection of object categories with a large number of object exemplars per category. To maintain engagement and encourage deep processing, participants performed a continuous recognition task during scanning. Each participant was presented with a set of 9,000 unique images plus 1,000 images shared with the others. Images were presented in 4-second trials (> 9,000 trials in total per subject). Each image was presented three times over 30 to 40 scan sessions on different days. Eight subjects participated in that study.

### Memorability scoring

The 73,000 natural images were rated with a memorability score using ViTMem. The score ranges from 0 (low memorability) to 1 (high memorability). Figure 1 presents the distribution of memorability scores.

**Figure 1.**
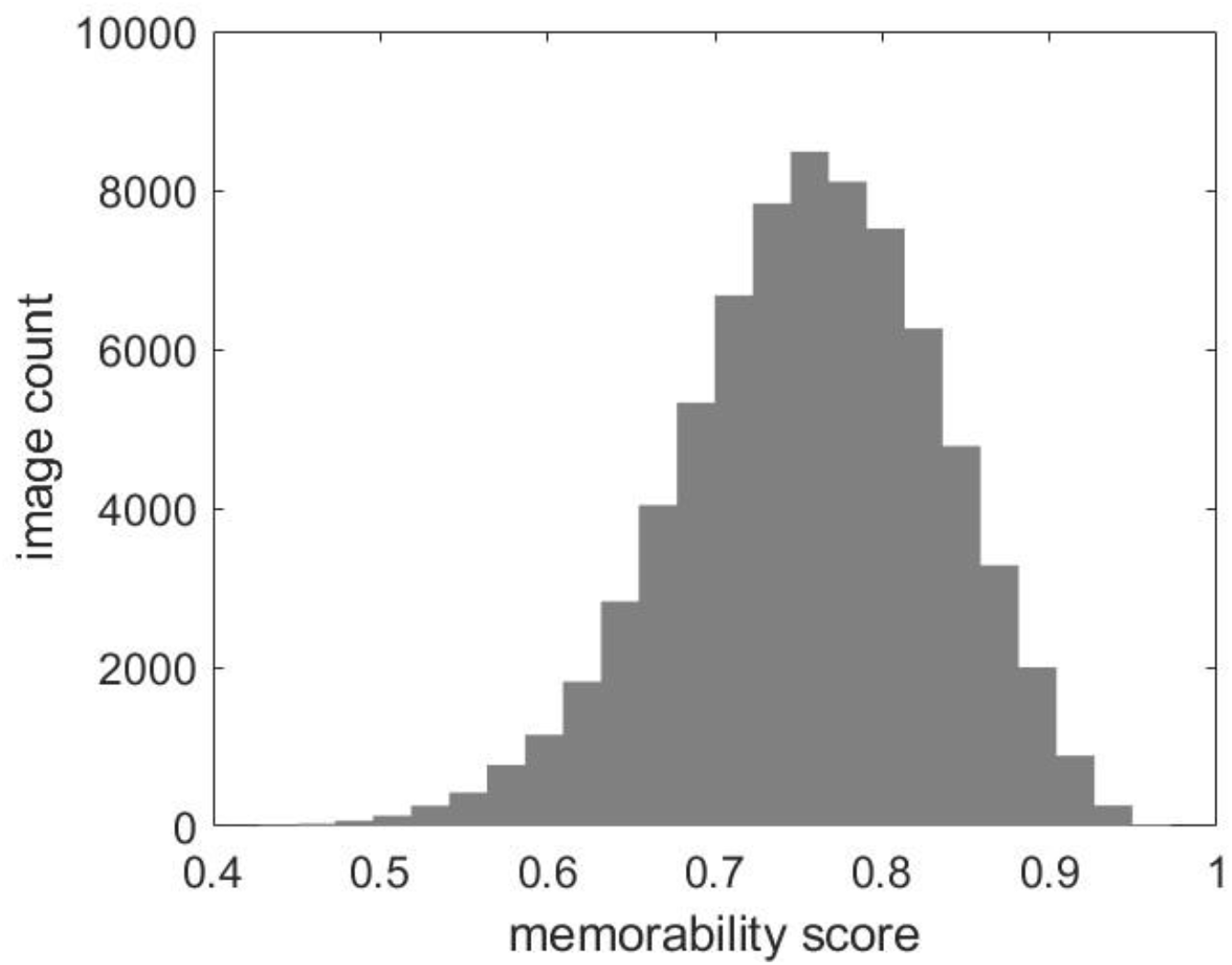
Histogram of memorability values of all images of the dataset.

### fMRI analysis

The beta values associated with the presentation of each image in each session were used for the current analyses. To avoid confounding effects of repetition and recognition effects (Bainbridge & Rissman 2018), we only included the presentation of each image for the first time, with a total of 10,000 trials per subject. We first performed first-level regression models including the memorability scores as regressors, obtaining one map of brain activation associated with increasing levels of memorability. Contrast images were smoothed with a Gaussian kernel (FWHM = 4 mm^3^) and submitted to a second-level analysis to evaluate the group effects of memorability scores. The analyses were run using SPM toolbox for fMRI implemented in MATLAB. Anatomical labels of coordinates were obtained with the Talairach toolbox (Lancaster et al. 2000).

## Results

We obtained group-level brain maps of brain activation to image presentation as a function of their memorability scores (Figure 2, Table 1). We found that higher memorability scores were associated with increased activation in middle occipital areas and fusiform gyrus. We also observed a relationship in medial temporal lobe areas, the hippocampus, and a set of subcortical regions such as amygdala and thalamus. These results are consistent with previous studies, which have implicated the ventral visual pathway and medial temporal lobe in processing image memorability, while primary visual areas typically do not show such activity. In addition, we found a significant effect on inferior, middle and supplementary frontal areas, as well as in parietal regions, including the superior and inferior parietal cortex.

**Figure 2.**
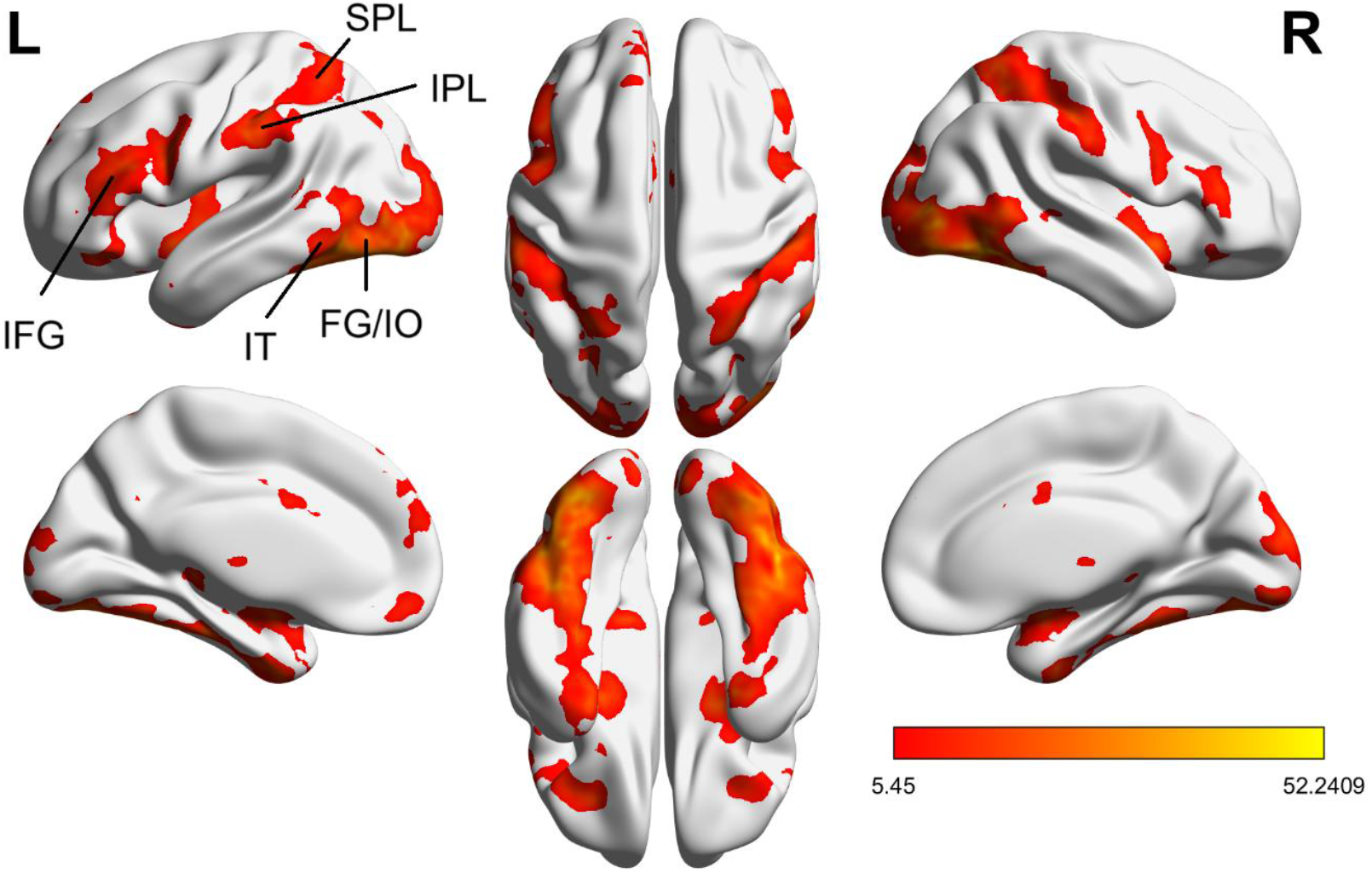
Brain regions where activation to images correlated with memorability score. IFG: inferior frontal gyrus; SPL: superior parietal lobe; IPL: inferior parietal lobe; IT: inferior temporal region; FG: fusiform gyrus; IO: inferior occipital area.

**Table1.**
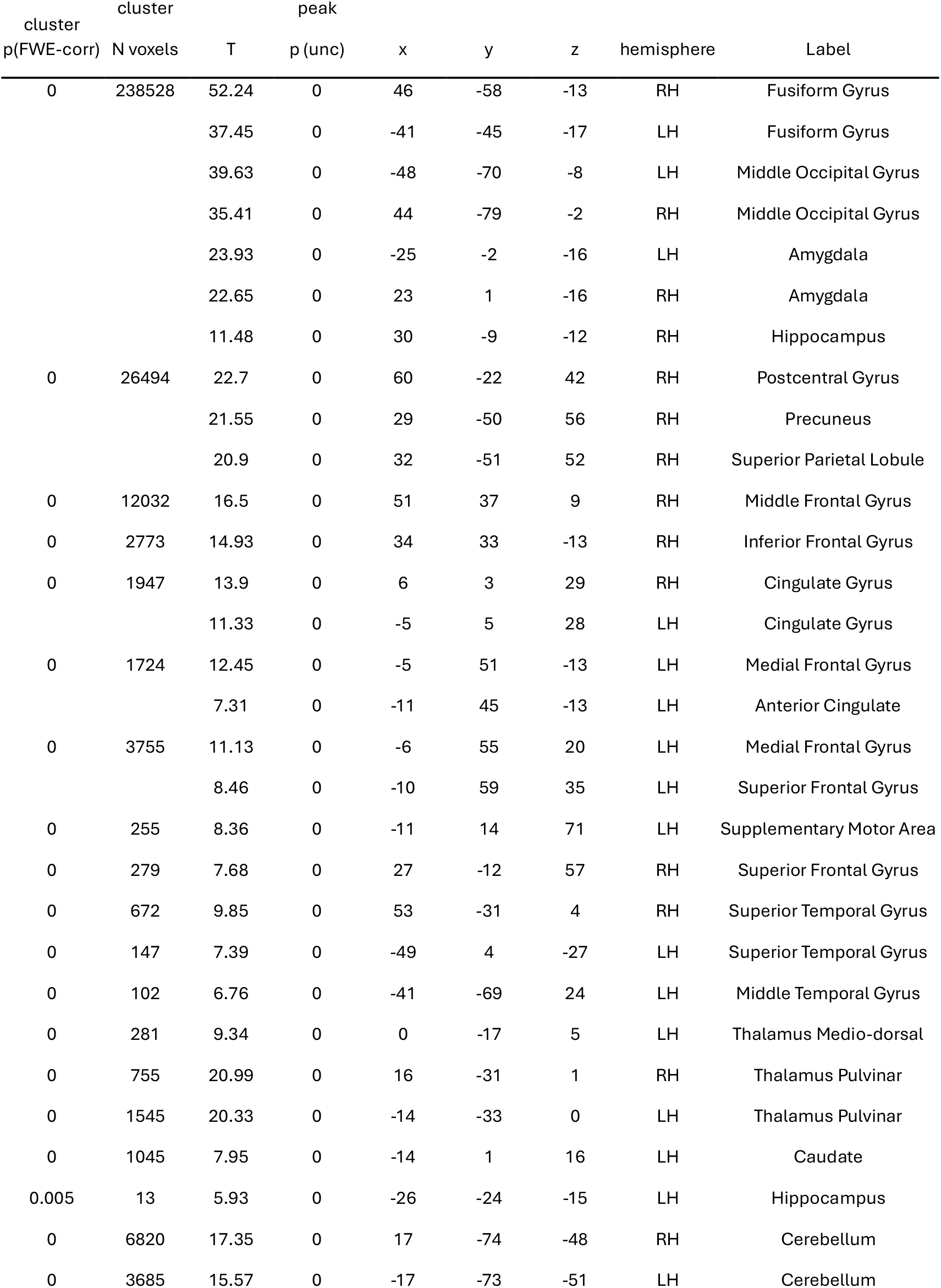
Coordinates with significant effects of the memorability score on region activation associated with image presentation. LH: left hemisphere; RH: right hemisphere; PFC: prefrontal cortex; Inf: inferior; Mid: middle; Sup: superior; Post: posterior; unc.: uncorrected; AAL: automatic anatomical labelling; BA: Brodmann area.

## Discussion

Our findings reveal that higher memorability scores are linked to increased activation to image presentation in inferior and middle occipital areas, as well as medial temporal lobe regions, including amygdala and hippocampus. These results align with previous studies implicating the ventral visual pathway and medial temporal lobe in processing image memorability, while primary visual areas show little involvement. Additionally, we observed significant effects in bilateral frontal and parietal regions, suggesting a broader network contributing to image memorability beyond regions implicated with image memorability in previous studies. These studies had a limited sample size (N = 15 to 18, one session each) and examined a comparatively small number of images across memorability levels (from N = 78 to N ~ 700). Here we used a highly sampled dataset of >240 sessions of high-field fMRI, and >70,000 images, which significantly increased our power to detect smaller effects, as could be the case in associative areas. The fronto-parietal findings suggest that higher areas may be implicated in modulating the degree to which they are later remembered, and may be thus an important part of the circuit mediating perception-memory processes. Furthermore, many of the fronto-parietal regions are associated with working memory, sensory-motor and premotor circuits (inferior frontal gyrus, supplementary motor area, and superior parietal lobule). A possibility is thus that memorability may modulate behavior through involvement of higher order regions. Some evidence suggests that ocular behavior is related to strength of memory formation (Popov & Staudigl, 2023). Future studies are necessary to assess the mechanisms and behavioral significance of graded image memorability.

Overall, our findings demonstrate that ANN-based memorability scoring using ViTMem aligns closely with previously reported neural correlates from subject-based image ratings. This consistency suggests that ViTMem provides a reliable and promising approach for investigating how fundamental visual stimulus properties influence brain activity and cognitive processing. More broadly, our study highlights the viability of combining ANNs trained to predict image memorability with large-scale fMRI datasets.

This approach offers a powerful framework for further exploring the neural mechanisms underlying image memorability, paving the way for more efficient and scalable investigations into perceptual and memory-related processes.

